# Post-transcriptional control by RNA-binding proteins links local synaptic translation to schizophrenia genetic risk

**DOI:** 10.64898/2026.02.27.708510

**Authors:** Kamilė Tamušauskaitė, Philippa M. Wells, Akshay Bhinge, Jonathan Mill, Nicholas E Clifton

**Affiliations:** Department of Clinical & Biomedical Sciences, University of Exeter Medical School, University of Exeter, Exeter, United Kingdom; Living Systems Institute, University of Exeter, Exeter, United Kingdom

**Keywords:** Schizophrenia, Local Translation, RNA-Binding Proteins, Synapse, Genome-wide association study

## Abstract

**Background:** Synaptic function is increasingly recognized as a core property of genes implicated in psychiatric disorders. Defining the specific synaptic molecular systems underlying genetic risk is critical step toward therapeutic advances. Synaptic processes rely on rapid protein production driven by local translation of mRNA in context-specific synapses. Synaptic mRNA metabolism, transport and local translation is regulated by RNA-binding proteins (RBPs). Here, we hypothesized that genetic risk converges on localised transcripts with synaptic function and aimed to identify RBP regulatory systems that capture this shared schizophrenia genetic risk.

**Methods:** We use recent human and mouse bulk and single-synapse transcriptomic and proteomic datasets to test for enrichment of schizophrenia genetic risk among mRNAs stratified by localization and synaptic function employing gene set association (MAGMA) and heritability enrichment (S-LDSC) analyses. Prioritized transcripts were further analyzed for RBP control through motif enrichment analysis (Transite) of the 3’UTRs of these transcripts. Candidate RBPs were then evaluated based on the strength of genetic association among their predicted binding targets.

**Results:** We demonstrate that genes encoding localised mRNAs with synaptic function show significantly greater genetic association than other synaptic genes. We identified a subset of RBPs, RBFOX1/2/3, CELF4, HNRNPR, and nELAVL, whose motifs are enriched in localised synaptic mRNAs and whose targets are enriched for schizophrenia risk variants. These RBPs are prioritized as candidate regulatory systems through which genetic risk may converge on the transport, splicing and translation of localised transcripts with synaptic function.

**Conclusions:** Our results highlight potential regulatory systems through which genetic variation influences synaptic mechanisms and provide a scalable framework for refining the link between genetic association and post-transcriptional regulation in neuropsychiatric disorders.

## Introduction

Schizophrenia ranks among the top 15 causes of disability worldwide^1^. Conventional antipsychotic drugs often lead to serious side effects^2^ and show variable efficacy^3^, reflecting the poorly understood pathoetiology of schizophrenia. It follows that progress in developing effective medication has been limited for over 70 years. Given that schizophrenia has a high heritability of 65-80%^4–6^, investigating genetic risk factors could provide insights into disease mechanisms.

Recent genomic studies have given insight into the complex polygenic architecture of schizophrenia, suggesting that the genetic risk converges on the structure and function of neuronal synapses^7–11^. To improve therapeutics and prevention strategies, understanding the specific mechanisms through which genetic variants confer risk is crucial. While excitatory and inhibitory neurons in the cortex and hippocampus are consistently implicated in schizophrenia^12,13^, each neuron can form multiple different types of synaptic connection, including both excitatory and inhibitory^14^. Synapses may independently undergo activity-driven changes, giving rise to distinct molecular signatures^15^. This underlies the need to explore the relative contribution to schizophrenia risk from different synaptic subtypes.

Neurons can synthesise synaptic proteins either within the soma or locally in remote compartments, such as dendrites, and the axon. The local translation of gene transcripts enables rapid protein production, supporting activity-dependent synaptic events essential for developmental processes such as synaptogenesis and neurite outgrowth and mature synaptic transmission and plasticity^16–18^, which have all been implicated in schizophrenia risk through genetic and functional studies^19–21^. The hypothesis that risk variants influencing synaptic function might be concentrated in locally translated genes was recently explored by Clifton et al.^22^, identifying schizophrenia association enriched in localised synaptic transcripts from mouse cortical synaptoneurosomes and the hippocampal CA1 dendritic translatome. The applicability of these findings to human synaptic biology, however, was unexplored and no study has yet determined the relative risk contributed by synaptic subtypes originating from distinct cell types. Moreover, while transcript transport, metabolism and translation are regulated by RNA-binding proteins (RBPs), it is yet to be determined whether the observed genetic association with schizophrenia in these localised synaptic transcripts could be mechanistically linked to RBP-mediated processes.

In the present study, using published human and mouse single-synapse transcriptomics and proteomics datasets in conjunction with the latest case-control genotype data, we explored the relative contribution to schizophrenia genetic risk in synaptically localised transcripts and proteins across a range of synapse subtypes, cell types and brain regions and determined the relationships between the genetic associations in the local synaptic translatome and RBP binding.

## Methods

### Gene sets

#### Synaptic gene ontology

Synaptic functional annotations were obtained from expert-curated Synaptic Gene Ontologies and annotations consortium (SynGO)^23^. In total, 1602 unique human gene IDs were included (SynGO release 1.2 aka “20231201”).

#### Synaptic transcriptome

Data containing transcripts localised to individual neurites, primarily composed of synaptosomes, and nuclear transcripts from human adult hippocampal excitatory cornu ammonis (CA) and dentate gyrus (DG) neurons, inhibitory hippocampal neurons and excitatory prefrontal cortex (PFC) neurons were obtained from Niu et al. 2023^24^. The synapse-enriched transcripts were defined by Niu et al.^24^ as those enriched in synaptosomes in comparison to the nucleus with a false discovery rate (FDR) < 0.05 and abs(log_2_(fold-change (FC)) > log_2_(1.3). The matched single-nucleus profiles from the same neuronal subtypes were used for downstream analysis to control for neuron-type-specific gene expression that is not related to synaptic localisation. Three new gene-sets were created by intersecting synaptic functional annotations from SynGO^23^ with transcriptomes of synaptosomes from Niu et al. 2023^24^: localised transcripts with synaptic function (L-Syn), localised transcripts without synaptic function (L-NonSyn) and non-localised transcripts with synaptic function (N-Syn) (Figure 1A).

**Figure 1.**
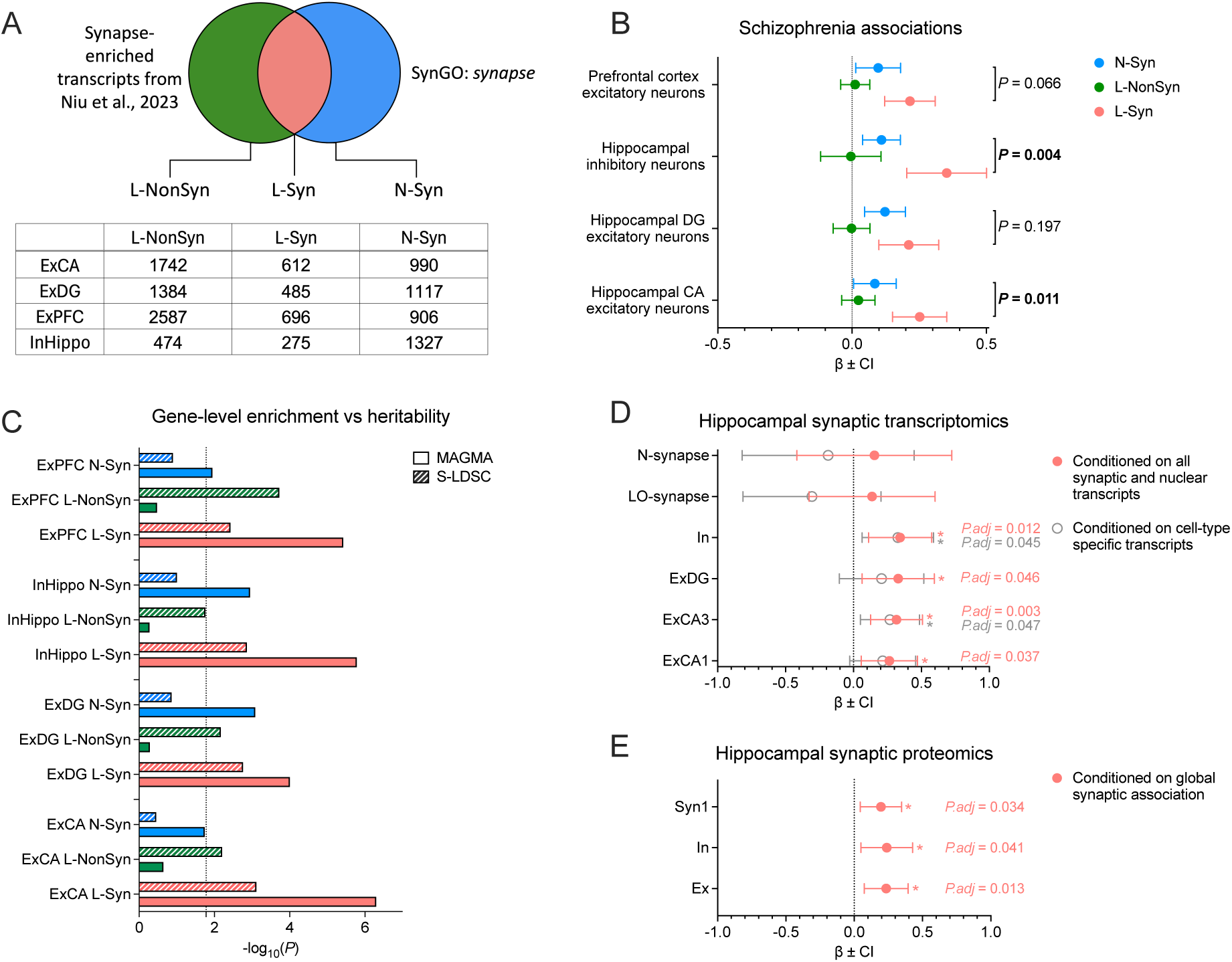
(A) Overlap between synapse-localised mRNAs and SynGO synaptic annotations. L-Syn describes mRNA enriched in the local transcriptome and annotated to SynGO:synapse (red). L-NonSyn refers to mRNA enriched in the local transcriptome but not annotated to SynGO:synapse (green). N-Syn represents mRNA annotated to SynGO:synapse but not enriched in the local transcriptome (blue). The table represent a number of genes in each gene set derived from the intersection, followed by gene symbol conversion to human NCBI Entrez Gene IDs. **(B)** Common variant enrichment for schizophrenia across gene groups defined by mRNA localisation and synaptic function. *P*-values reflect z-test comparisons of effect sizes from MAGMA competitive gene set analysis between L-Syn and N-Syn. **(C)** MAGMA competitive gene set association (solid bars) comparison with heritability enrichment from S-LDSC (hatched bars) in gene sets defined by synaptic function and mRNA localisation. Bars demonstrate the significance of association −log_10_(*P*) in MAGMA and S-LDSC. The dashed line indicates the threshold for significance after adjusting for multiple testing using the Bonferroni method. **(D)** Common variant enrichment for schizophrenia across transcript groups defined by L-Syn within human hippocampal synapse subtypes. Red colour represent gene sets conditioned on all genes expressed in total nuclei and synapses across hippocampal subregions in MAGMA competitive gene set analysis. Grey circles show common variant enrichment for schizophrenia after conditioning on transcripts specifically expressed in pyramidal CA1 neurons, interneurons and medium spiny neurons, demonstrating effects independent of cell-type enrichment. *P*-values adjusted using Bonferroni correction. **(E)** Common variant enrichment for schizophrenia across protein groups defined L-Syn within inhibitory (In), excitatory (Ex) and broad (Syn1) mouse hippocampal synapse subtypes. *P*-values adjusted using Bonferroni correction. Displayed values represent effect sizes (*β*) ± confidence intervals (CI) from MAGMA competitive gene set analysis. ExCA: cornu ammonis excitatory synapses; ExDG: dentate gyrus excitatory synapses; ExPFC: prefrontal cortex excitatory synapses; InHippo/In: hippocampal inhibitory synapses; LO-synapse: low RNA abundance synapses; N-synapse: nascent transcript enriched synapses.

Additionally, subtype-specific synaptic transcriptomes from Niu et al. 2023^24^ were extracted, including excitatory and inhibitory synapse subtypes from human hippocampus (CA1 excitatory (ExCA1), CA3 excitatory (ExCA3), DG excitatory (ExDG), inhibitory (In), low RNA abundance synapse (LO-synapse) and nascent transcript enriched synapse (N-synapse). The transcripts from these subtypes were intersected with SynGO to focus exclusively on localised transcripts annotated with synaptic function.

#### Synaptic proteome

Single-synapse hippocampal proteomes of adult mice from Camk2a+, Gad2+ and Syn1 Cre-driver lines were acquired from Van Oostrum et al. 2023^25^. In this study, the synapse-type-specific proteomes were intersected with SynGO to prioritise proteins with synaptic functions.

#### Synaptic transcriptome of hippocampal regions and strata

Whole-tissue and localised synaptic transcriptomes from broad hippocampal subregions (CA1, CA3, and DG) and four CA1 strata (stratum oriens (SO), stratum pyramidale (SP), and stratum radiatum (SR) and stratum lacunosum-moleculare (SLM)) were obtained from Kaulich et al. 2025^26^. When indicated, these datasets were intersected with SynGO to identify L-Syn transcripts.

### GWAS summary statistics

Genome-wide association study (GWAS) summary statistic data from individuals with schizophrenia were obtained from a recent meta-analysis of 74,776 cases and 101,023 control individuals of European, East Asian, Latino and African American ancestry^7^. SNPs were filtered to only include those with high imputation quality (imputation INFO score ≥ 0.8) and minor allele frequency (MAF) ≥ 0.01. GWAS summary statistics of the standing height trait were obtained from Pan-UKB listed under phenotype code 50, which included 419,596 European ancestry individuals^27^. Low confidence variants flagged by the dataset were removed and SNPs with MAF ≤ 0.01 were filtered out to exclude very rare variants.

### Gene-set enrichment analysis in MAGMA

Gene-set enrichment analysis for common genetic variants was performed using multiple regression models in Multi-marker Analysis of GenoMic Annotation (MAGMA) v1.10^28^. SNPs were mapped to genes using the Ensembl GRCh37 genome and population data from the European Phase 3 1000 Genomes Project was used to account for linkage disequilibrium. Both data files were acquired as auxiliary files from the MAGMA website (https://cncr.nl/research/magma/). SNPs were annotated to genes with a 35kb upstream and 10kb downstream window around gene boundaries to include proximal regulatory regions. The *SNP-wise Mean* model was used to determine gene-wide *P*-values using GWAS SNP *P*-values. One-tailed competitive gene-set analyses were undertaken to estimate the strength of genetic associations in gene sets of interest while adjusting for potential confounders such as gene length, SNP density, and differences in SNP sample sizes. Human single-synapse transcriptomic datasets from the PFC and hippocampus were analysed using conditional MAGMA gene-set analysis to account for known associations between brain expressed genes and psychiatric phenotypes. This was undertaken by conditioning on all synapse– and nucleus-enriched transcripts from any cell type and all SynGO genes. For hippocampal single-synapse subtype analysis, conditioning was restricted to synapse– and nucleus-enriched transcripts expressed across hippocampal regions, together with SynGO genes to determine independent associations beyond those shared by genes expressed broadly in the constituent neural tissue. To further assess independent associations beyond the cell type enrichment^12^, hippocampal subtype analyses were covaried on continuous gene-level cell type specificity scores relating to pyramidal CA1 neurons, interneurons and medium spiny neurons obtained from Skene et al. 2018^9^, which were previously implicated in schizophrenia. Single-synapse hippocampal proteomes were conditioned on globally expressed synaptic genes together with SynGO genes. Statistical comparison of effect sizes was performed using z-test of β values. *P*-values were adjusted using Bonferroni correction to control for testing of multiple gene-sets.

### Developmental expression-association analysis

MAGMA interaction analysis was used to determine the relationship between GWAS association and developmental gene expression within a gene set. Relative expression scores for each gene across developmental stages were obtained from a previously published study^29^. These scores were calculated using RNA-seq transcriptomic data from and hippocampal post-mortem prenatal (14 to 22 post-conception weeks; n = 57) and adult (18 to 96 years; n = 460) brain tissue samples, sourced from the BrainSeq Phase II^30^. MAGMA interaction analyses were two-tailed and *P*-values were Bonferroni adjusted for 10 developmental stages assessed.

### Dataset preparation for Stratified LD Score Regression

Protein coding transcript and 3’UTR coordinates were defined using GENCODE v46 lifted over to GRCh37. Gene coordinates were defined by collapsing all gene features and taking the earliest start and latest end coordinate per gene. Consolidated 3’UTR regions were generated by collapsing overlapping 3’UTR fragments. Gene and transcript 3’UTR annotations to RBP targets were established from overlapping coordinates. Unannotated RBP target regions were removed.

### Stratified LD Score Regression (S-LDSC)

The enrichment for schizophrenia heritability was evaluated using S-LDSC^31^. The reference genome was obtained from the 1000 Genomes Project (European). Heritability partitioning was performed by regressing PGC3 schizophrenia summary statistics (INFO ≥ 0.8; MAF ≥ 0.01) on LD scores and SNPs were annotated to each gene using a 10 kb window around each gene^32^. The 97 functional baseline annotations were used to control for confounders such as functional elements, conservation scores, chromatin marks, which are already associated with heritability^33^. One-sided *P*-values were calculated from coefficient z-score to determine if SNPs in RBP annotations explain more heritability than would be expected by chance.

### RBP motif gene set enrichment analysis

To identify which RBPs are likely to have a role in regulating localised synaptic transcripts, RBPs motif enrichment analysis in the 3’UTRs of these localised transcripts was performed using a previously published method^34^. Protein coding transcript 3’UTR sequences were extracted from human gencode v46. For each localised synaptic transcript, overlapping 3’UTR fragments were collapsed by gene and strand to generate consolidated 3’UTR regions. Motif position weight matrices (PWM) were obtained from the Transite database^35^. The universalmotif R package was used for motif scanning in the 3’ UTR sequences with a relative log-odds threshold of 0.8. Enrichment was determined by undertaking gene set enrichment analysis (GSEA) for each motif using the fgsea package, and *P*-values were adjusted using the Benjamini-Hochberg method to control the FDR for multiple testing (alpha = 0.05).

### Annotation, filtering and mapping pipeline for RBP data using POSTAR3 peaks

ECLIP-derived targets of selected RBPs were retrieved from POSTAR3^36^. For eCLIP datasets originating from K562 and HepG2 human cell lines (ENCODE), peaks from both cell lines were combined and mapped to overlapping protein coding genes using the human genome assembly GRCh38 (Ensembl release 102). Peaks matched to multiple genes and sites with no overlapping genes were excluded from downstream analysis.

### Manual extraction and processing pipeline for RBP targets from published studies

We curated RBP targets from published human and mouse brain or neuronal CLIP datasets (Supplementary Table 1). Unannotated peaks were mapped to protein-coding murine genes on GRCm37 before conversion to human homologs. Peaks matched to multiple genes and sites with no overlapping genes were excluded from downstream analysis.

### Gene ID conversion

Mouse and human gene identifiers were converted to human NCBI Entrez Gene IDs using Bioconductor package biomaRt^37^. To ensure consistent mapping, archived Ensembl 104 release BioMart data (GRCh38/mm10) was used. Only one-to-one mappings were retained and genes without a valid Entrez ID were excluded from downstream analyses, retaining ∼95% of the starting gene list after Entrez ID conversion.

### RBP target stratification by binding confidence for enrichment analysis

RBP targets were ranked by binding confidence provided by POSTAR3, defined using input-normalised peak *P*-values for each gene. Where multiple peaks were annotated to a gene, their *P*-values were combined using Simes test using R package mppa (version 1.0). In manually extracted RBP target datasets, *P*-values from multiple peaks associated with the same gene were combined using Simes’ test; when *P*-values were not available, the mean rank across peaks per gene was used instead. The top ranked up to 800 RBP target genes were used in primary analyses. Additional bins of up to 400 genes, by rank, were used to evaluate the change in gene set association with decreasing RBP binding confidence.

## Results

### Local synaptic transcripts carry the strongest enrichment for schizophrenia association

Competitive tests adjusting for background association from each cell type revealed that in synapses from human hippocampal excitatory CA (ExCA) neurons, schizophrenia associations were enriched in genes encoding localised transcripts with synaptic functions (L-Syn) (β = 0.252, Bonferroni adjusted *P (P.adj)* = 1.50 × 10^−6^). A comparison of effect sizes confirmed a stronger enrichment in L-Syn compared to non-localised mRNAs with synaptic functions (N-Syn) (z = 2.547, *P* = 0.011) and compared to localised mRNAs without synaptic functions (L-NonSyn) (z = 3.774, *P* = 1.60 × 10^−4^). In synapses from hippocampal inhibitory neurons (InHippo) and human DG excitatory hippocampal neurons (ExDG), schizophrenia associations were enriched in genes encoding both L-Syn (InHippo: β = 0.352, *P.adj =* 4.94 × 10^−6^; ExDG: β = 0.211, *P.adj =* 2.97 × 10^−4^) and N-Syn (InHippo: β = 0.110, *P.adj =* 0.003; ExDG: β = 0.122, *P.adj =* 0.002). Again, InHippo transcripts had a much stronger genetic association in L-Syn compared to N-Syn (z = 2.898, *P* = 0.004) or L-NonSyn (z = 3.760, *P* = 1.70 × 10^−4^). However, in ExDG, the effect size among L-Syn and N-Syn genes did not significantly vary by localization (z = 1.290, *P* = 0.197), suggesting that synaptic transcripts in excitatory DG are enriched for schizophrenia association regardless of their localisation site. In human excitatory PFC synapses, schizophrenia associations were also enriched in both L-Syn and N-Syn (L-Syn: β = 0.215, *P.adj =* 1.13 × 10^−5^; N-Syn: β = 0.097, *P.adj =* 0.034) although neither was stronger than the other (z = 1.840, *P* = 0.066). Among localised transcripts, L-Syn carried greater association than L-NonSyn (z = 3.663, *P* = 2.50 × 10^−4^) (Figure 1B). The overlap between L-Syn datasets is shown in Supplementary Figure 1A.

In each cell type, both L-Syn and L-NonSyn demonstrated enrichment of schizophrenia SNP-heritability, indicating that schizophrenia risk variants are concentrated within genes whose transcripts localize to synapses. However, only genes in L-Syn demonstrated both gene-level association (MAGMA) and significant heritability enrichment (S-LDSC). These results provide genetic evidence that disruptions of core synaptic molecular processes, rather than generic synaptic transcript localization, represent a causal component of schizophrenia biology (Figure 1C).

### Schizophrenia association enriched in excitatory and inhibitory synapses in synaptic transcriptome of human hippocampus

Having established that genetic association with schizophrenia is enriched in L-Syn genes, we examined how this association was distributed among synapse subtypes taken from single-synapse transcriptomes of human hippocampus.

We tested for independent genetic associations in single-synapse transcriptomes beyond broadly expressed neural and synaptic genes. We observed an enrichment for schizophrenia genetic association in L-Syn from excitatory CA1, CA3 and DG hippocampal synapses (ExCA1: β = 0.264, *P.adj =* 0.037; ExCA3: β = 0.317, *P.adj =* 0.003; ExDG: β = 0.329, *P.adj =* 0.046), and inhibitory hippocampal synapses (β = 0.343, *P.adj =* 0.012), indicating that the enrichment in these subtypes exceeds cell-wide transcript localisation and synapse-related background association (Figure 1D).

Additional conditional analyses demonstrated that the enrichment for schizophrenia association in L-Syn CA3 synapses and hippocampal inhibitory synapses was independent of cell type-specific gene expression previously associated with schizophrenia^12^ (ExCA3: β = 0.268, *P.adj =* 0.047; InHippo: β = 0.326, *P.adj =* 0.045) (Figure 1D).

### Schizophrenia association enrichment in synaptic proteomes

We sought to determine whether synapse-specific genetic associations observed at the transcriptome level follow a similar pattern at the proteomic level, independent of the site of protein synthesis. Using single-synapse proteomes from the mouse hippocampus and restricting analysis to SynGO-annotated proteins while adjusting for global synaptic association, we found that proteins enriched in either hippocampal excitatory or inhibitory synapses were significantly enriched for association with schizophrenia (Syn1: β = 0.196, *P.adj =* 0.034; In: β = 0.238, *P.adj =* 0.041; Ex: β = 0.234, *P.adj =* 0.013) (Figure 1E). Thus, the comparable enrichment between excitatory and inhibitory L-Syn datasets identified at the transcript level in humans is partially reflected in the mouse proteome (Supplementary Figure 1B).

### Schizophrenia association enrichment in transcriptomes of hippocampal regions and strata

Following the identification of synapse subtype-specific transcriptomic associations with schizophrenia, we aimed to dissect their relative regional and laminar contributions across hippocampal subregions (DG, CA3 and CA1) and strata of CA1 (SP, SO, SR and SLM). We performed a stepwise analysis, firstly, we assessed all transcripts to identify overall regional and laminar enrichment, then intersected these with SynGO-annotated synaptic genes to determine if the signal was driven by genes with known synaptic functions. These analyses were repeated for bulk and synaptosomal transcriptomes.

While controlling for overall hippocampal gene expression, we found that transcripts broadly enriched in any of the three hippocampal subregions showed significant enrichment for schizophrenia common risk variants (CA1: β = 0.111, *P =* 1.00 × 10^−6^; CA3: β = 0.058, *P =* 0.005; DG: β = 0.106, *P =* 1.76 × 10^−6^). Within CA1 strata, only SP (β = 0.162 and *P =* 2.99 × 10^−9^) and SR (β = 0.130 and *P =* 0.010) layers exhibited significant associations. Notably, transcripts with synaptic function from all three hippocampal subregions were significantly enriched for schizophrenia-associated common variants, with the strongest enrichment observed in CA1 (CA1: β = 0.363, *P =* 4.34 × 10^−11^; CA3: β = 0.252, *P =* 1.12 × 10^−7^; DG: β = 0.185, *P =* 0.001). Among the layers of CA1, transcripts with synaptic functions in SO (β = 0.448, *P =* 0.048) and SP (β = 0.386, *P =* 1.75 × 10^−10^) showed significant enrichment for schizophrenia-linked variants (Figure 2).

**Figure 2.**
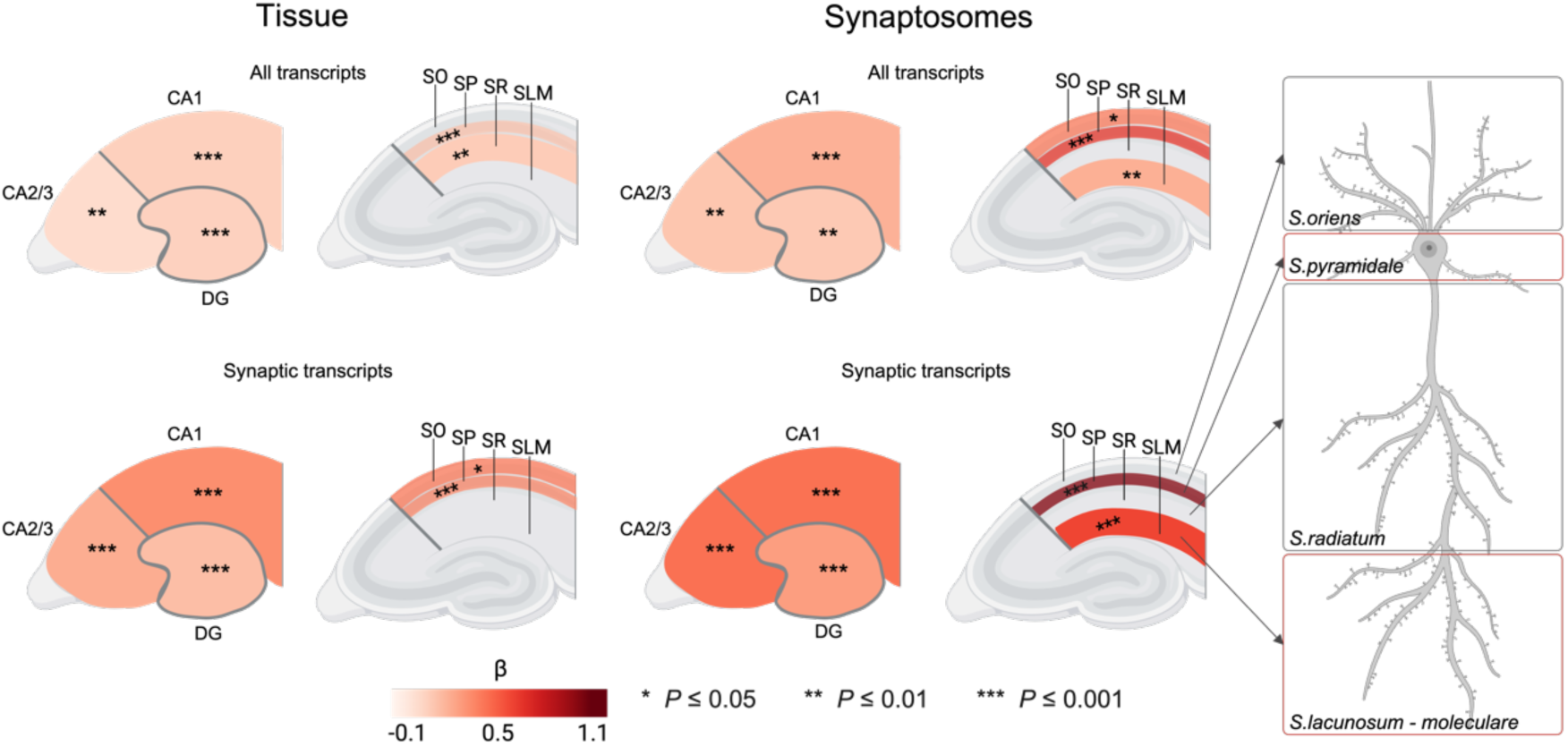
Schizophrenia genetic association across hippocampal regions and strata in whole-tissue and synaptosome transcriptomes. The schematic illustration on the left shows analyses of gene sets defined by regional transcript enrichment, including all regionally enriched transcripts across hippocampal regions and strata (top left) and SynGO-annotated synaptic transcripts that are regionally enriched (bottom left). The visual representation on the right shows analogous analyses restricted to transcripts enriched in synaptosomes across hippocampal regions and strata (top right) and L-Syn within these regions and strata (bottom right). Asterisks denote raw *P*-values and the colour bar indicates effect sizes (β), where darker colours represent stronger genetic association in MAGMA competitive gene set analysis. Figure created with BioRender.com. CA: cornu ammonis; DG: dentate gyrus; SLM: stratum lacunosum-moleculare; SO: stratum oriens; SP: stratum pyramidale; SR: stratum radiatum.

Next, we focused on transcriptomes from synaptosomal compartments to refine the regional and laminar associations observed in bulk tissue. Similarly to bulk tissue, transcripts localised to synapses showed significant enrichment for schizophrenia in all three hippocampal subregions compared to background expression (CA1: β = 0.234, *P =* 1.04 × 10^−6^; CA3: β = 0.156, *P =* 0.002; DG: β = 0.141, *P =* 0.003). The common variant association became even stronger when restricting to transcripts with known synaptic functions (CA1: β = 0.468, *P =* 6.99 × 10^−7^; CA3: β = 0.470, *P =* 1.23 × 10^−6^; DG: β = 0.306, *P =* 3.57 × 10^−4^). The L-Syn associations in CA1 and CA3 were greater than N-Syn (CA1: z = 2.882, *P* = 0.004; CA3: z = 2.820, *P* = 0.005). In DG, the effect size among L-Syn and N-Syn did not significantly vary by localization (z = 1.196, *P* = 0.232). Within the layers of CA1, transcripts localised to SO, SP and SLM synapses were significantly associated (SO: β = 0.440, *P =* 0.016; SP: β = 0.663, *P =* 2.42 × 10^−5^; SLM: β = 0.238, *P =* 0.004), although the association in the SO layer was not prevalent when restricting to L-Syn (SP: β = 1.022, *P =* 4.13 × 10^−6^; SLM: β = 0.620, *P =* 1.44 × 10^−4^). In SP and SLM layers, the association within L-Syn was greater than N-Syn (SP: z = 3.586, *P* = 3.36 × 10^−4^; SLM: z = 2.458, *P* = 0.014) (Figure 2).

### Schizophrenia association among L-Syn is not related to developmental expression

So far, we have established that L-Syn of human CA excitatory neurons and human hippocampal inhibitory neurons show a strong association with schizophrenia. Next, focusing on the hippocampus, we explored whether gene expression during certain developmental periods is related to enrichment for common variant association for schizophrenia in L-Syn. In L-Syn of human ExCA and InHippo neurons, both at the level of cell type or synapse subtype, no significant interactions were observed between relative developmental expression and genetic association with schizophrenia at any stage, following Bonferroni correction (Supplementary Figure 1C).

### Developmental expression profile of inhibitory and excitatory synaptic proteins in the hippocampus

We explored whether gene expression during specific developmental periods is related to the schizophrenia common variant association attributed to single-synapse protein expression. In hippocampal excitatory synapses, we observed a significant positive interaction between gene expression during early infancy and genetic association with schizophrenia (β = 0.098, *P.adj =* 0.015). No further significant interactions were observed in any the datasets tested (Supplementary Figure 1C).

### RNA-binding protein motif analysis of L-Syn 3’UTRs

We next aimed to investigate potential post-transcriptional mechanisms characterising the schizophrenia association in L-Syn. In 3’UTR of L-Syn from human CA excitatory neurons and inhibitory hippocampal neurons, we found 100 and 14 enriched RBP motifs, which are bound by 85 and 16 unique RBPs, respectively (FDR < 0.05). All 14 motifs enriched in 3’UTRs of L-Syn in InHippo synapses overlapped with those enriched in L-Syn excitatory CA synapses (Figure 3A).

**Figure 3.**
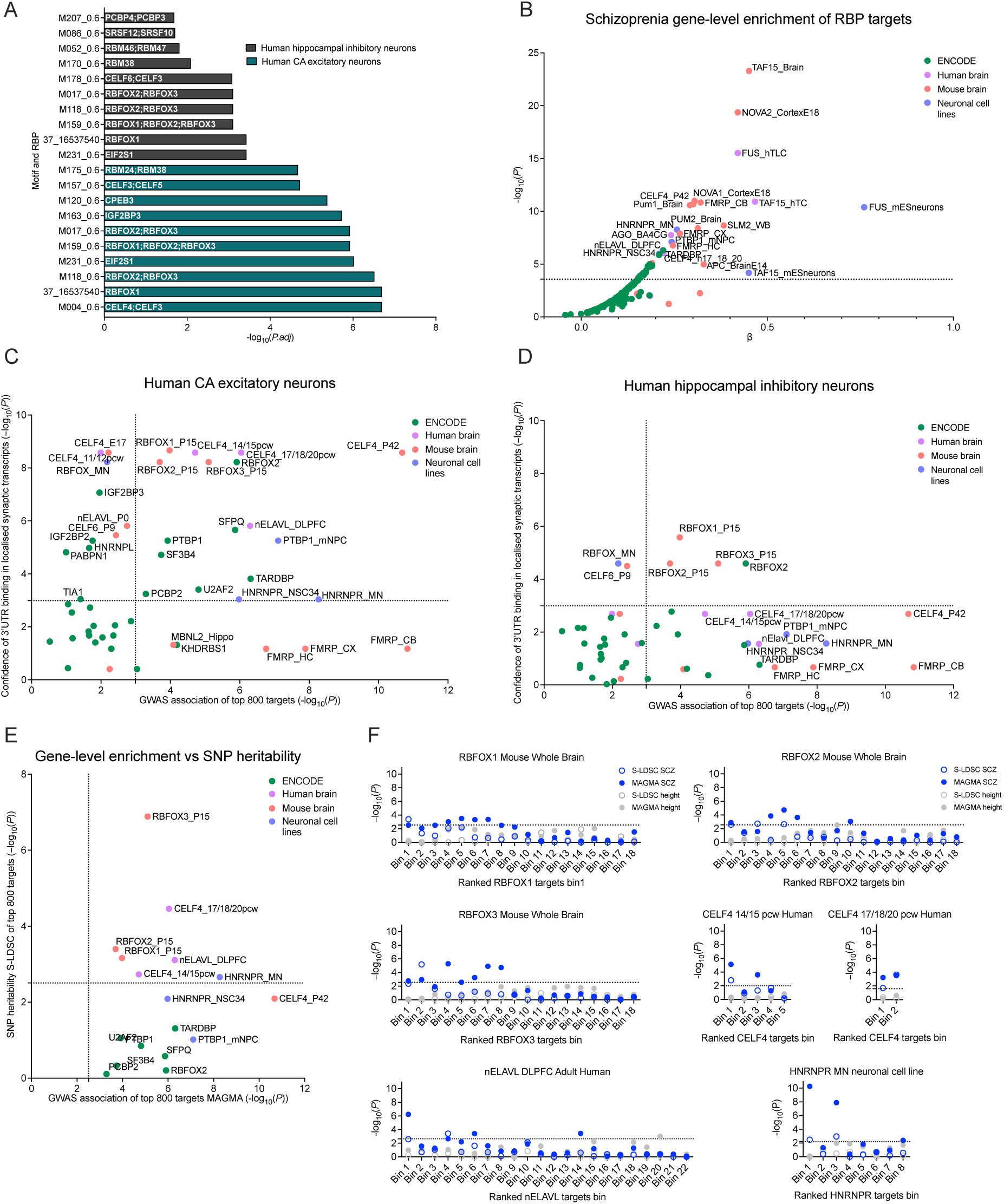
(A) Top 10 RBP motifs enriched in 3’UTRs of L-Syn in human hippocampal inhibitory and CA excitatory neurons. Values represent –log_10_ FDR-adjusted *P*-values (FDR < 0.05) from GSEA for each motif and respective RBP. **(B)** Common variant enrichment for schizophrenia across up to 800 top binding targets of RBPs derived from ENCODE K562 and HepG2 cell lines, neuronal cell lines and human and mouse brain tissues. The dashed line indicates the threshold for significance after adjusting MAGMA gene-set enrichment *P*-values using the Bonferroni method. **(C)** and **(D)** Common genetic association with schizophrenia compared with L-Syn 3’UTR motif enrichment across RBP target sets. Top motif enrichment (–log_10_ *P*-values) for each RBP was compared with MAGMA gene-set enrichment *P*-values for up to 800 top RBP targets. Dotted lines represent thresholds for significance after adjusting for multiple testing using the Bonferroni method. **(E)** Comparison of gene set enrichment and SNP heritability −log_10_(*P*) values for RBP target sets identified by MAGMA. The dotted line represents the threshold for significance after adjusting enrichment *P*-values using the Bonferroni method for 16 gene sets. **(F)** The relationship between enrichment for schizophrenia (blue) or height (grey) genetic association and RBP binding confidence for six prioritised RBPs. Genes were ranked by RBP-mRNA CLIP binding scores and divided into bins of 400 genes with the first bin containing the highest CLIP scores. The number of bins reflects the number of genes for which ranking information was available from the primary dataset. Each bin was tested for association using MAGMA competitive get set analysis and SNP heritability enrichment using S-LDSC. Datapoints correspond to the −log_10_(*P*) of each test. The dashed line indicates the threshold for significance after adjusting for the number of bins using the Bonferroni method.

### Schizophrenia associations of RBP targets identified in K562, HepG2 and neuronal cell lines and brain tissue

Next, we sought to assess the schizophrenia association of RBP target genes. Focusing on ENCODE eCLIP datasets from two cell lines, K562 and HepG2, we identified several RBPs (e.g., TARDBP, RBFOX2, and SFPQ) whose top targets exhibited a modest yet statistically significant positive association with schizophrenia risk (β = 0.2 – 0.3). We then examined RBP targets from mouse and human brain tissues and neuronal cell lines. We identified numerous additional RBP target sets (e.g., TAF15, NOVA1/2, FUS, CELF4, and FMRP) with strongly significant association with schizophrenia risk (β = 0.3 – 0.8) (Figure 3B) as well as further evidence for association in targets of a subset of RBPs highlighted from ENCODE (e.g. RBFOX2 and PTBP1).

### Common variant enrichment for schizophrenia across targets of RBPs positively enriched within the 3ʹUTRs of localised synaptic transcripts

We integrated the data above to identify RBP targets with strong 3’UTR binding to L-Syn and exhibiting enrichment for schizophrenia-associated variants. We report 12 RBPs, which appear to regulate L-Syn that may be particularly important in conferring schizophrenia risk (Figure 3C and 3D). We next applied S-LDSC to examine whether variants mapping to these 12 RBP target genes explained a significant proportion of schizophrenia SNP heritability. Targets of six of the 12 RBPs showed significant enrichment in both MAGMA and S-LDSC analyses, providing convergent evidence across gene-level and variant-level analyses (Figure 3E). Among these prioritized RBPs, top binding targets of splicing regulators Rbfox1/2/3 in whole mouse brain tissue showed consistent enrichment for schizophrenia risk. Moreover, while translation regulator CELF4 targets vary from early fetal development to late midfetal stage in the neocortex^38^, only the binding targets defined between 14 to 20 postconceptional weeks in human neocortex were enriched for schizophrenia risk. In contrast, we observed no SNP heritability enrichment among the top Celf4 targets in 6-week-old mouse whole brain. Targets of post-transcriptional regulators nELAVL in human adult dorsolateral prefrontal cortex and Hnrnpr in motoneurons were also enriched for schizophrenia risk.

### The relationship between RBP binding confidence and association with schizophrenia

We further analyzed the prioritized RBPs (RBFOX1/2/3, CELF4, nELAVL and HNRNPR) by assessing whether schizophrenia risk enrichment scales with RBP binding confidence across successive binding bins of 400 genes using both MAGMA and S-LDSC. In both analyses, bins comprising genes with higher RBP binding confidence showed stronger enrichment for schizophrenia risk. Unlike schizophrenia, height-associated variants did not show enrichment in bins of genes with higher RBP binding confidence (Figure 3F).

### Developmental expression profile of prioritized RBP targets and schizophrenia association

We undertook MAGMA interaction analysis to determine whether gene expression of prioritized RBP targets across developmental stages is related to their enrichment for common variant association for schizophrenia. The enrichment of prioritized RBP targets did not correlate with specific developmental period in the hippocampus, suggesting that their association with schizophrenia is not driven by changes in expression over development (Supplementary Figure 1C).

## Discussion

Local protein synthesis is crucial for synaptic plasticity, and its dysregulation has been implicated in schizophrenia^22^, proposing that perturbations in the transport, stability and local translation of specific synaptic mRNAs may be a key pathway in disease aetiology. We demonstrate schizophrenia risk enrichment among a subset of genes with synaptic functions whose mRNAs are trafficked to synapses for local translation and identify key RBPs, RBFOX1/2/3, CELF4, HNRNPR, and nELAVL, that may contribute to the regulation of these transcripts.

Previous findings suggested that schizophrenia risk attributed to L-Syn is common to excitatory and inhibitory neurons^22^. Our analysis of single-synapse transcriptomes supports this notion as we observed broad genetic associations across each of the major hippocampal synapse subtypes. This observation was mirrored in single-synapse proteomic data, showing that the strength of genetic association was unaffected by excitatory or inhibitory status. While the proteomic data reflect transcripts translated both in the soma and the synapse, the synaptic transcriptome contains candidates for local translation, placing an importance on activity-dependent excitatory and inhibitory processes in schizophrenia pathology. Notably, the enrichment of schizophrenia-associated variants in L-Syn over N-Syn was region-specific. In human hippocampal excitatory CA and inhibitory synapses, but not in excitatory DG and PFC synapses, the L-Syn signal was significantly stronger than N-Syn. This pattern was consistent in mouse hippocampal synaptosomes, where L-Syn transcript association was significantly greater than N-Syn in CA1 and CA3, but not in DG. Within CA1, this enrichment in L-Syn over N-Syn was further refined to synapses of the SP and SLM layers, which contain inhibitory synapses onto pyramidal cell bodies and distal excitatory inputs (e.g., from entorhinal cortex), respectively. This suggests that the genetic risk contribution of L-Syn is particularly relevant to the microcircuitry of the hippocampal CA subfields and local inhibitory networks, whereas synaptic risk in the DG and PFC may be characterised by mechanisms independent of transcript localisation.

Our analysis identified RBPs whose motifs are enriched in the 3’UTRs of L-Syn transcripts from inhibitory and excitatory synapses of hippocampus. Several of these, including RBFOX1, RBFOX2, ELAVL1, and FXR2, have previously been reported to regulate cortical synaptic transcriptomes through 3’UTR binding^39^. While we observed a greater number of RBP motifs enriched in excitatory over inhibitory L-Syn transcript 3’UTRs in hippocampal synapses, all motifs enriched in the inhibitory synapses overlapped with those from CA excitatory synapses, consistent with observations from cortical synaptic transcriptomes^39^. Among the RBPs associated with either synapse type, we identified several, such as RBFOX1/2/3, CELF4, HNRNPR, and nELAVL, whose target genes show substantial enrichment for genetic association with schizophrenia. Furthermore, the confidence scores for binding by these RBPs correlated with the strength of schizophrenia association among their targets, suggesting that genes under stronger regulatory influence from these RBPs are more likely to harbour risk variants. Many RBP target sets identified in neuronal tissues displayed a stronger association with schizophrenia risk than those derived from ENCODE cell lines, underscoring the importance of tissue specificity in RBP function and relevance to psychiatric disorders. For numerous RBPs, neuronal targets remain incompletely defined, forming a gap that likely obscures additional regulators of L-Syn transcripts relevant to schizophrenia risk.

With regards to their biological roles, the RBFOX family proteins, whose targets and the genes themselves are implicated in schizophrenia and other psychiatric disorders^40–42^, may regulate L-Syn transcripts undergoing nuclear pre-processing prior to localisation to synapses^43^ or through context-dependent functions beyond splicing. HNRNPR, which has been linked to neurodevelopmental disorders^44,45^, likely orchestrates the transport and local translation initiation of L-Syn^46–48^. CELF4, whose targets and the gene itself is strongly associated with various psychiatric conditions^40,49–53^, may maintain these L-Syn mRNAs in a translationally repressed state until locally activated^38,54^. Finally, nELAVL proteins may stabilise these L-Syn mRNAs preventing degradation and/or promote their local translation in response to synaptic signals^55,56^. While direct evidence for nELAVLs in schizophrenia is limited, the family members and their target genes are implicated in ASD and genes with synaptic function^52,57^, suggesting that the role of nELAVL in schizophrenia risk may be underexplored. Although RBFOX1/2/3, CELF4, HNRNPR, and nELAVL each regulate diverse RNA populations across developmental time points and subcellular contexts, they may form one or more disease-relevant regulatory modules controlling L-Syn transcripts under their combinatorial actions, rather than as isolated regulators of their target transcripts^58^. In turn, this may perturb synaptic activity in a time-, cell type-, region– and context-specific manner, thereby contributing to schizophrenia pathogenesis.

Notably, the genetic association of L-Syn was not correlated with age-specific developmental gene expression in the hippocampus. This suggests that the genetic risk captured by L-Syn transcripts may be conferred via synaptic functions that are maintained across several developmental stages, or that are not sufficiently captured by bulk tissue transcriptomics data. Instead, we observed an isolated signal linking genetic association among excitatory synaptic proteins to gene expression during early infancy. Schizophrenia association within prioritised RBP targets also did not correlate with expression across developmental stages, although to examine this further, more work needs to be done to characterise changes in RBP target genes over development. For instance, we observed temporal specificity for CELF4 targets, which were enriched in mid-fetal neocortex but not in early fetal neocortex, indicating that certain RBP-associated effects on risk could be restricted to specific developmental periods.

This study has several limitations. Firstly, we were unable to directly compare the 3’UTR of excitatory and inhibitory L-Syn transcripts due to a lack of isoform-specific data. Consequently, we could not determine if the greater number of RBP motifs observed in transcripts localised to excitatory synapses was due to longer 3’UTR and greater cis-regulatory complexity as previously suggested^39,59,60^. Secondly, our RBP motif enrichment analysis was conducted on L-Syn transcripts from adult tissue only. Since both RBP targets and L-Syn expression may change across development, the RBPs regulating L-Syn during development could differ from those identified here and we cannot yet assess whether schizophrenia risk variants preferentially disrupt RBP programs in early development. Indeed, gene expression during early developmental periods has previously been linked to schizophrenia risk^61–63^. Our analyses do not test whether the schizophrenia associated SNPs affect RBP binding. However, previous evidence suggests that variants affecting RBP-mRNA interactions contribute significantly to the heritability of psychiatric disorders^64^. During RBP prioritization, technical constraints of the Transite motif database precluded enrichment analysis for a subset of RBPS, including TAF15 and FUS, whose targets demonstrated strong schizophrenia associations. FUS is known to mediate mRNA transport to dendrites^65^ and regulate stability of numerous synaptic mRNAs^39,66–68^. TAF15 is involved in alternative splicing of synaptic mRNAs^69^. Thus, the enrichment of schizophrenia associations among FUS and TAF15 targets, together with prior evidence of their involvement in synaptic regulation, suggests that FUS and TAF15 may regulate L-Syn and may play a role in conferring schizophrenia genetic risk. Finally, while NOVA2 targets from fetal brain were enriched for schizophrenia associations, we detected no NOVA2 motif enrichment in adult hippocampal L-Syn transcripts. This discrepancy likely reflects the sharp decline of NOVA2 expression after early development^70,71^, underscoring the temporal specificity of RBP regulation. These limitations collectively highlight the need for future studies to map RBP targets across development in disease-relevant tissues and to utilize isoform-specific, single-synapse transcriptomics and proteomics for a complete understanding of L-Syn transcript regulation.

Together, these results highlight potential regulatory systems through which genetic variation influences synaptic mechanisms and provide a scalable framework for refining the link between genetic association and post-transcriptional regulation in neuropsychiatric disorders.

## Acknowledgements

We thank Prof Erin Schuman group for early access to key datasets and the Psychiatric Genomics Consortium for sharing the genomics datasets used in this study. This project utilised the University of Exeter High-Performance Computing (HPC) facility funded by the UK Medical Research Council (MRC), Clinical Research Infrastructure Initiative (MR/M008924/1). This work was funded by grant MR/W017156/1 from the MRC to NEC.

## Bibliography

1. Vos T, Abajobir AA, Abate KH, et al. Global, regional, and national incidence, prevalence, and years lived with disability for 328 diseases and injuries for 195 countries, 1990–2016: a systematic analysis for the Global Burden of Disease Study 2016. The Lancet. 2017;390(10100):1211–1259. doi:10.1016/s0140-6736(17)32154-2

2. De Berardis D, Rapini G, Olivieri L, et al. Safety of antipsychotics for the treatment of schizophrenia: a focus on the adverse effects of clozapine. Ther Adv Drug Saf. May 2018;9(5):237–256. doi:10.1177/2042098618756261

3. Lieberman JA, Stroup TS, McEvoy JP, et al. Effectiveness of antipsychotic drugs in patients with chronic schizophrenia. N Engl J Med. Sep 22 2005;353(12):1209–23. doi:10.1056/NEJMoa051688

4. Cardno AG, Gottesman, II. Twin studies of schizophrenia: from bow-and-arrow concordances to star wars Mx and functional genomics. Am J Med Genet. Spring 2000;97(1):12–7.

5. Hilker R, Helenius D, Fagerlund B, et al. Heritability of Schizophrenia and Schizophrenia Spectrum Based on the Nationwide Danish Twin Register. Biol Psychiatry. Mar 15 2018;83(6):492–498. doi:10.1016/j.biopsych.2017.08.017

6. Sullivan PF, Kendler KS, Neale MC. Schizophrenia as a Complex Trait: Evidence From a Meta-analysis of Twin Studies. Archives of General Psychiatry. 2003;60(12):1187–1192. doi:10.1001/archpsyc.60.12.1187

7. Trubetskoy V, Pardinas AF, Qi T, et al. Mapping genomic loci implicates genes and synaptic biology in schizophrenia. Nature. Apr 2022;604(7906):502–508. doi:10.1038/s41586-022-04434-5

8. Singh T, Poterba T, Curtis D, et al. Rare coding variants in ten genes confer substantial risk for schizophrenia. Nature. Apr 2022;604(7906):509–516. doi:10.1038/s41586-022-04556-w

9. Nakamura T, Takata A. The molecular pathology of schizophrenia: an overview of existing knowledge and new directions for future research. Molecular Psychiatry. 2023;28(5):1868–1889. doi:10.1038/s41380-023-02005-2

10. Fromer M, Pocklington AJ, Kavanagh DH, et al. De novo mutations in schizophrenia implicate synaptic networks. Nature. Feb 13 2014;506(7487):179–84. doi:10.1038/nature12929

11. Hall J, Bray NJ. Schizophrenia Genomics: Convergence on Synaptic Development, Adult Synaptic Plasticity, or Both? Biological Psychiatry. 2022;91(8):709–717. doi:10.1016/j.biopsych.2021.10.018

12. Skene NG, Bryois J, Bakken TE, et al. Genetic identification of brain cell types underlying schizophrenia. Nature Genetics. 2018;50(6):825–833. doi:10.1038/s41588-018-0129-5

13. Ruzicka WB, Mohammadi S, Fullard JF, et al. Single-cell multi-cohort dissection of the schizophrenia transcriptome. Science. 2024;384(6698)doi:10.1126/science.adg5136

14. Megías M, Emri Z, Freund TF, Gulyás AI. Total number and distribution of inhibitory and excitatory synapses on hippocampal CA1 pyramidal cells. Neuroscience. 2001;102(3):527–40. doi:10.1016/s0306-4522(00)00496-6

15. Cizeron M, Qiu Z, Koniaris B, et al. A brainwide atlas of synapses across the mouse life span. Science. 2020;369(6501):270–275. doi:10.1126/science.aba3163

16. Holt Christine E, Schuman Erin M. The Central Dogma Decentralized: New Perspectives on RNA Function and Local Translation in Neurons. Neuron. 2013/10/30/ 2013;80(3):648–657. 10.1016/j.neuron.2013.10.036

17. Holt CE, Martin KC, Schuman EM. Local translation in neurons: visualization and function. Nature Structural & Molecular Biology. 2019/07/01 2019;26(7):557–566. doi:10.1038/s41594-019-0263-5

18. Rajgor D, Welle TM, Smith KR. The Coordination of Local Translation, Membranous Organelle Trafficking, and Synaptic Plasticity in Neurons. Front Cell Dev Biol. 2021;9:711446. doi:10.3389/fcell.2021.711446

19. Hall J, Trent S, Thomas KL, O’Donovan MC, Owen MJ. Genetic Risk for Schizophrenia: Convergence on Synaptic Pathways Involved in Plasticity. Biological Psychiatry. 2015/01/01/ 2015;77(1):52–58. 10.1016/j.biopsych.2014.07.011

20. Pardinas AF, Holmans P, Pocklington AJ, et al. Common schizophrenia alleles are enriched in mutation-intolerant genes and in regions under strong background selection. Nat Genet. Mar 2018;50(3):381–389. doi:10.1038/s41588-018-0059-2

21. Berdenis van Berlekom A, Muflihah CH, Snijders GJLJ, et al. Synapse Pathology in Schizophrenia: A Meta-analysis of Postsynaptic Elements in Postmortem Brain Studies. Schizophrenia Bulletin. 2019;46(2):374–386. doi:10.1093/schbul/sbz060

22. Clifton NE, Lin JQ, Holt CE, O’Donovan MC, Mill J. Enrichment of the Local Synaptic Translatome for Genetic Risk Associated With Schizophrenia and Autism Spectrum Disorder. Biological Psychiatry. 2024;95(9):888–895. doi:10.1016/j.biopsych.2023.12.006

23. Koopmans F, van Nierop P, Andres-Alonso M, et al. SynGO: An Evidence-Based, Expert-Curated Knowledge Base for the Synapse. Neuron. Jul 17 2019;103(2):217–234.e4. doi:10.1016/j.neuron.2019.05.002

24. Niu M, Cao W, Wang Y, et al. Droplet-based transcriptome profiling of individual synapses. Nature Biotechnology. 2023/01/16 2023;doi:10.1038/s41587-022-01635-1

25. Van Oostrum M, Blok TM, Giandomenico SL, et al. The proteomic landscape of synaptic diversity across brain regions and cell types. Cell. 2023;186(24):5411–5427.e23. doi:10.1016/j.cell.2023.09.028

26. Kaulich E, Waselenchuk Q, Fürst N, et al. An integrated transcriptomic and proteomic map of the mouse hippocampus at synaptic resolution. Nat Commun. Aug 26 2025;16(1):7942. doi:10.1038/s41467-025-63119-5

27. Karczewski KJ, Gupta R, Kanai M, et al. Pan-UK Biobank genome-wide association analyses enhance discovery and resolution of ancestry-enriched effects. Nature Genetics. 2025/10/01 2025;57(10):2408–2417. doi:10.1038/s41588-025-02335-7

28. de Leeuw CA, Mooij JM, Heskes T, Posthuma D. MAGMA: generalized gene-set analysis of GWAS data. PLoS Comput Biol. Apr 2015;11(4):e1004219. doi:10.1371/journal.pcbi.1004219

29. Clifton NE, Collado-Torres L, Burke EE, et al. Developmental Profile of Psychiatric Risk Associated With Voltage-Gated Cation Channel Activity. Biol Psychiatry. Sep 15 2021;90(6):399–408. doi:10.1016/j.biopsych.2021.03.009

30. Collado-Torres L, Burke EE, Peterson A, et al. Regional Heterogeneity in Gene Expression, Regulation, and Coherence in the Frontal Cortex and Hippocampus across Development and Schizophrenia. Neuron. 2019;103(2):203–216.e8. doi:10.1016/j.neuron.2019.05.013

31. Finucane HK, Bulik-Sullivan B, Gusev A, et al. Partitioning heritability by functional annotation using genome-wide association summary statistics. Nature Genetics. 2015/11/01 2015;47(11):1228–1235. doi:10.1038/ng.3404

32. Clifton NE, Schulmann A, Holmans PA, O’Donovan MC, Vawter MP. The relationship between case-control differential gene expression from brain tissue and genetic associations in schizophrenia. Am J Med Genet B Neuropsychiatr Genet. Jul-Sep 2023;192(5-6):85–92. doi:10.1002/ajmg.b.32931

33. Gazal S, Finucane HK, Furlotte NA, et al. Linkage disequilibrium–dependent architecture of human complex traits shows action of negative selection. Nature Genetics. 2017/10/01 2017;49(10):1421–1427. doi:10.1038/ng.3954

34. Hawkins S, Mondaini A, Namboori SC, et al. ePRINT: exonuclease assisted mapping of protein-RNA interactions. Genome Biology. 2024;25(1)doi:10.1186/s13059-024-03271-1

35. Krismer K, Bird MA, Varmeh S, et al. Transite: A Computational Motif-Based Analysis Platform That Identifies RNA-Binding Proteins Modulating Changes in Gene Expression. Cell Reports. 2020;32(8):108064. doi:10.1016/j.celrep.2020.108064

36. Zhao W, Zhang S, Zhu Y, et al. POSTAR3: an updated platform for exploring post-transcriptional regulation coordinated by RNA-binding proteins. Nucleic Acids Research. 2022;50(D1):D287–D294. doi:10.1093/nar/gkab702

37. Durinck S, Spellman PT, Birney E, Huber W. Mapping identifiers for the integration of genomic datasets with the R/Bioconductor package biomaRt. Nature Protocols. 2009;4(8):1184–1191. doi:10.1038/nprot.2009.97

38. Salamon I, Park Y, Miškić T, et al. Celf4 controls mRNA translation underlying synaptic development in the prenatal mammalian neocortex. Nature Communications. 2023;14(1)doi:10.1038/s41467-023-41730-8

39. Juengling M, Mosbacher J, Perez J, et al. Brain-wide synaptosome profiling reveals localized mRNAs that diversify synapses. bioRxiv. 2025:2025.12.07.692806. doi:10.64898/2025.12.07.692806

40. Genovese G, Fromer M, Stahl EA, et al. Increased burden of ultra-rare protein-altering variants among 4,877 individuals with schizophrenia. Nature Neuroscience. 2016/11/01 2016;19(11):1433–1441. doi:10.1038/nn.4402

41. De Rubeis S, He X, Goldberg AP, et al. Synaptic, transcriptional and chromatin genes disrupted in autism. Nature. 2014/11/01 2014;515(7526):209–215. doi:10.1038/nature13772

42. Bigdeli TB, Chatzinakos C, Bendl J, et al. Biological insights into schizophrenia from ancestrally diverse populations. Nature. 2026/01/21 2026;doi:10.1038/s41586-025-10000-6

43. Weyn-Vanhentenryck SM, Mele A, Yan Q, et al. HITS-CLIP and integrative modeling define the Rbfox splicing-regulatory network linked to brain development and autism. Cell Rep. Mar 27 2014;6(6):1139–1152. doi:10.1016/j.celrep.2014.02.005

44. Gillentine MA, Wang T, Hoekzema K, et al. Rare deleterious mutations of HNRNP genes result in shared neurodevelopmental disorders. Genome Med. Apr 19 2021;13(1):63. doi:10.1186/s13073-021-00870-6

45. Duijkers FA, McDonald A, Janssens GE, et al. HNRNPR Variants that Impair Homeobox Gene Expression Drive Developmental Disorders in Humans. Am J Hum Genet. Jun 6 2019;104(6):1040–1059. doi:10.1016/j.ajhg.2019.03.024

46. Glinka M, Herrmann T, Funk N, et al. The heterogeneous nuclear ribonucleoprotein-R is necessary for axonal β-actin mRNA translocation in spinal motor neurons. Human Molecular Genetics. 2010;19(10):1951–1966. doi:10.1093/hmg/ddq073

47. Briese M, Saal-Bauernschubert L, Ji C, et al. hnRNP R and its main interactor, the noncoding RNA 7SK, coregulate the axonal transcriptome of motoneurons. Proc Natl Acad Sci U S A. Mar 20 2018;115(12):E2859–e2868. doi:10.1073/pnas.1721670115

48. Zare A, Salehi S, Bader J, et al. hnRNP R promotes O-GlcNAcylation of eIF4G and facilitates axonal protein synthesis. Nature Communications. 2024/08/28 2024;15(1):7430. doi:10.1038/s41467-024-51678-y

49. Howard DM, Adams MJ, Clarke T-K, et al. Genome-wide meta-analysis of depression identifies 102 independent variants and highlights the importance of the prefrontal brain regions. Nature Neuroscience. 2019/03/01 2019;22(3):343–352. doi:10.1038/s41593-018-0326-7

50. Levey DF, Stein MB, Wendt FR, et al. Bi-ancestral depression GWAS in the Million Veteran Program and meta-analysis in >1.2 million individuals highlight new therapeutic directions. Nature Neuroscience. 2021/07/01 2021;24(7):954–963. doi:10.1038/s41593-021-00860-2

51. Meng X, Navoly G, Giannakopoulou O, et al. Multi-ancestry genome-wide association study of major depression aids locus discovery, fine mapping, gene prioritization and causal inference. Nature Genetics. 2024/02/01 2024;56(2):222–233. doi:10.1038/s41588-023-01596-4

52. Satterstrom FK, Kosmicki JA, Wang J, et al. Large-Scale Exome Sequencing Study Implicates Both Developmental and Functional Changes in the Neurobiology of Autism. Cell. Feb 6 2020;180(3):568–584.e23. doi:10.1016/j.cell.2019.12.036

53. Barone R, Fichera M, De Grandi M, et al. Familial 18q12.2 deletion supports the role of RNA-binding protein CELF4 in autism spectrum disorders. Am J Med Genet A. Jun 2017;173(6):1649–1655. doi:10.1002/ajmg.a.38205

54. Wagnon JL, Briese M, Sun W, et al. CELF4 regulates translation and local abundance of a vast set of mRNAs, including genes associated with regulation of synaptic function. PLoS Genet. 2012;8(11):e1003067. doi:10.1371/journal.pgen.1003067

55. Mulligan MR, Bicknell LS. The molecular genetics of nELAVL in brain development and disease. European Journal of Human Genetics. 2023/11/01 2023;31(11):1209–1217. doi:10.1038/s41431-023-01456-z

56. Mirisis AA, Carew TJ. The ELAV family of RNA-binding proteins in synaptic plasticity and long-term memory. Neurobiology of Learning and Memory. 2019/05/01/ 2019;161:143–148. 10.1016/j.nlm.2019.04.007

57. Berto S, Usui N, Konopka G, Fogel BL. ELAVL2-regulated transcriptional and splicing networks in human neurons link neurodevelopment and autism. Hum Mol Genet. Jun 15 2016;25(12):2451–2464. doi:10.1093/hmg/ddw110

58. Khoroshkin M, Buyan A, Dodel M, et al. Systematic identification of post-transcriptional regulatory modules. Nature Communications. 2024/09/09 2024;15(1):7872. doi:10.1038/s41467-024-52215-7

59. Tushev G, Glock C, Heumüller M, Biever A, Jovanovic M, Schuman EM. Alternative 3′ UTRs Modify the Localization, Regulatory Potential, Stability, and Plasticity of mRNAs in Neuronal Compartments. Neuron. 2018;98(3):495–511.e6. doi:10.1016/j.neuron.2018.03.030

60. Bae B, Miura P. Emerging Roles for 3’ UTRs in Neurons. Int J Mol Sci. May 12 2020;21(10)doi:10.3390/ijms21103413

61. Gulsuner S, Walsh T, Watts Amanda C, et al. Spatial and Temporal Mapping of De Novo Mutations in Schizophrenia to a Fetal Prefrontal Cortical Network. Cell. 2013;154(3):518–529. doi:10.1016/j.cell.2013.06.049

62. Jaffe AE, Straub RE, Shin JH, et al. Developmental and genetic regulation of the human cortex transcriptome illuminate schizophrenia pathogenesis. Nature Neuroscience. 2018/08/01 2018;21(8):1117–1125. doi:10.1038/s41593-018-0197-y

63. Clifton NE, Hannon ES, Harwood JC, et al. Dynamic expression of genes associated with schizophrenia and bipolar disorder across development. Translational Psychiatry. 2019;9(1)doi:10.1038/s41398-019-0405-x

64. Park CY, Zhou J, Wong AK, et al. Genome-wide landscape of RNA-binding protein target site dysregulation reveals a major impact on psychiatric disorder risk. Nat Genet. Feb 2021;53(2):166–173. doi:10.1038/s41588-020-00761-3

65. Fujii R, Okabe S, Urushido T, et al. The RNA Binding Protein TLS Is Translocated to Dendritic Spines by mGluR5 Activation and Regulates Spine Morphology. Current Biology. 2005;15(6):587–593. doi:10.1016/j.cub.2005.01.058

66. Sahadevan S, Hembach KM, Tantardini E, et al. Synaptic FUS accumulation triggers early misregulation of synaptic RNAs in a mouse model of ALS. Nature Communications. 2021/05/21 2021;12(1):3027. doi:10.1038/s41467-021-23188-8

67. Udagawa T, Fujioka Y, Tanaka M, et al. FUS regulates AMPA receptor function and FTLD/ALS-associated behaviour via GluA1 mRNA stabilization. Nature Communications. 2015/05/13 2015;6(1):7098. doi:10.1038/ncomms8098

68. Yokoi S, Udagawa T, Fujioka Y, et al. 3’UTR Length-Dependent Control of SynGAP Isoform α2 mRNA by FUS and ELAV-like Proteins Promotes Dendritic Spine Maturation and Cognitive Function. Cell Rep. Sep 26 2017;20(13):3071–3084. doi:10.1016/j.celrep.2017.08.100

69. Ibrahim F, Maragkakis M, Alexiou P, Maronski MA, Dichter MA, Mourelatos Z. Identification of in vivo, conserved, TAF15 RNA binding sites reveals the impact of TAF15 on the neuronal transcriptome. Cell Rep. Feb 21 2013;3(2):301–8. doi:10.1016/j.celrep.2013.01.021

70. Saito Y, Yuan Y, Zucker-Scharff I, et al. Differential NOVA2-Mediated Splicing in Excitatory and Inhibitory Neurons Regulates Cortical Development and Cerebellar Function. Neuron. 2019;101(4):707–720.e5. doi:10.1016/j.neuron.2018.12.019

71. Piton A. NOVA1/2 genes and alternative splicing in neurodevelopment. Current Opinion in Genetics & Development. 2025/08/01/ 2025;93:102373. 10.1016/j.gde.2025.102373

