## Supplementary Figure 1 for "Post-transcriptional control by RNA-binding proteins links local synaptic translation to schizophrenia genetic risk"

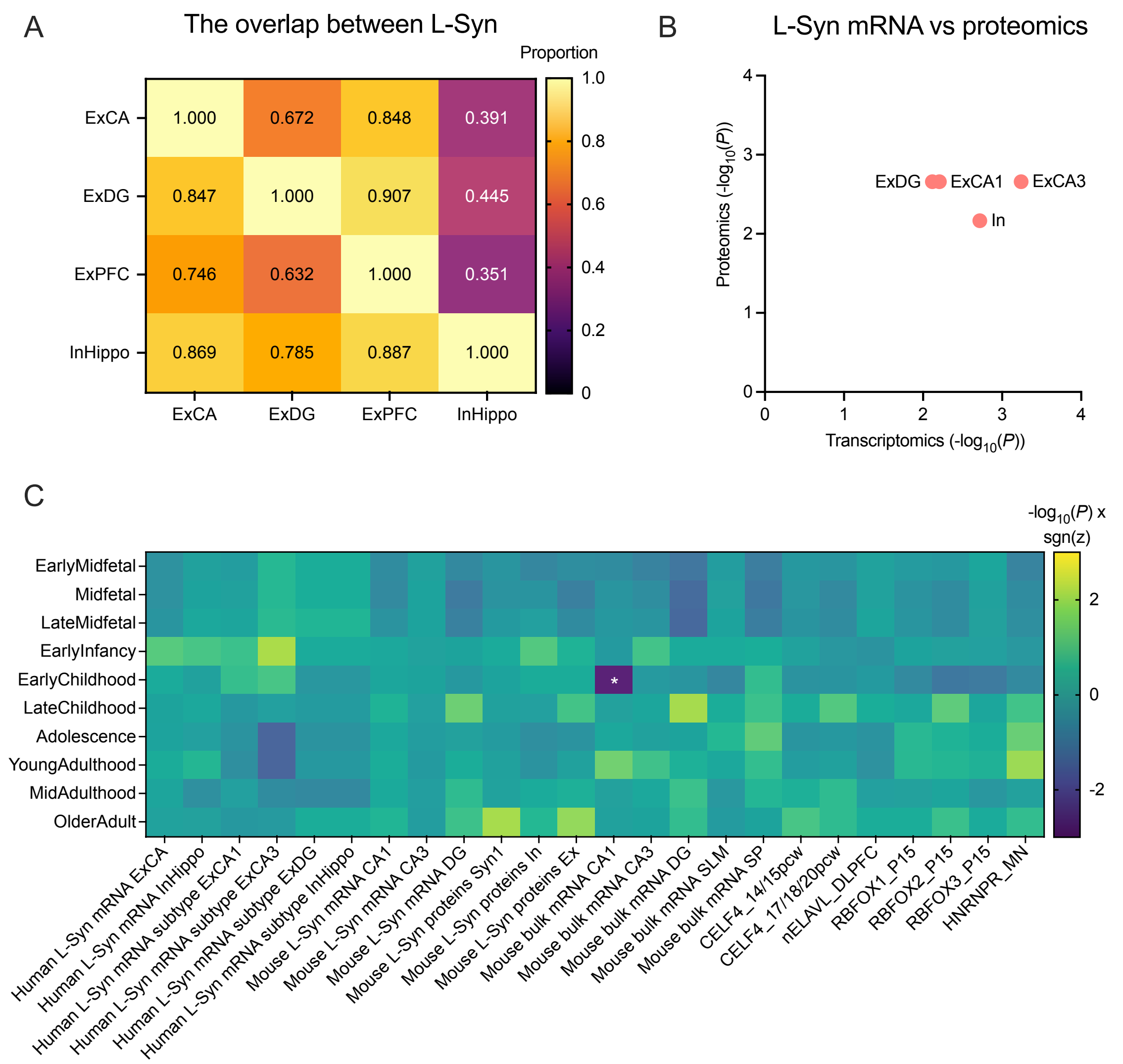


Supplementary Figure 1. **(A)** The overlap of L-Syn gene sets across human synaptic subtypes**.** The heatmap represents the proportional overlap ranging from 0 (no overlap) to 1 (identical sets) of L-Syn gene sets in each synaptic subtype (reference, rows) with L-Syn gene sets from other subtypes (columns) **(B)** Concordance of schizophrenia genetic association across matched hippocampal synapse subtypes between human L-Syn transcripts (x-axis) and mouse L-Syn proteome (y-axis). The plotted values demonstrate $-\log_{10}(P)$ derived from MAGMA competitive gene-set analyses after conditioning on respective global backgrounds. **(C)** MAGMA interaction analysis between the gene expression across developmental stages and hippocampal bulk and L-Syn transcripts and proteins, and prioritized RBP binding targets (up to 800 genes). Developmental hippocampal gene expression scores were obtained from BrainSeq Phase II. The heatmap colours represent $-\log_{10}(P) \times \mathrm{sgn}(z)$, where *P* is uncorrected following MAGMA interaction analysis, and the sign corresponds to the direction of the regression coefficient (z). The significance is considered after accounting for multiple testing across 10 developmental stages using the Bonferroni correction. ExCA: cornu ammonis excitatory synapses; ExDG: dentate gyrus excitatory synapses; ExPFC: prefrontal cortex excitatory synapses; InHippo/In: Hippocampal inhibitory synapses; L-Syn: localised transcripts with synaptic functions; MN: Motor neurons; P15: postnatal day 15; Pcw: postconceptual weeks; SLM: stratum lacunosum-moleculare; SP: stratum pyramidale.
